# Ex vivo drug testing in metastatic biopsies reveals patient-specific vulnerabilities to cancer targeting and immune activating drugs

**DOI:** 10.64898/2026.02.06.704037

**Authors:** Lukas Wöhrl, Durdam Das, Kathrin Weidele, Steffi Treitschke, Celine Baron, Dagmar Halbritter, Catherine Botteron, Florian Lüke, Nataša Stojanović Gužvić, Melanie Werner-Klein, Dennis Christoph Harrer, Tobias Pukrop, Julia Lanznaster, Thorsten Nitsch, Thomas Südhoff, Nina Fischer, Boris Kubuschok, Rainer Claus, Michaela Benz, Volker Bruns, Martin Hoffmann, Andrea Stutz, Christoph A. Klein, Christian Werno

## Abstract

Biomarker-guided therapies in oncology often fail to induce considerable responses in patients with advanced cancer. As a complementary approach, direct drug testing on individual patient samples is highly attractive yet is currently hampered by the lack of assays that combine (i) fast reporting, (ii) the ability to inform about immune-mediated responses, (iii) robust quantification, and (iv) scalability for parallel assessment of multiple drugs. Here, we introduce our patient-derived ex vivo drug response assay (PEDRA) that fulfills all these requirements. Using malignant pleural effusions (MPEs) from five non-small cell lung cancer (NSCLC) patients with detailed clinical treatment histories, we tested 52 guideline-recommended therapies and eight investigational antibody-drug conjugates (ADCs). In all patients, PEDRA identified treatment options that outperformed the therapies the patients had received. The results reflected clinical observations as well as expectations derived from mutational profiling and disease courses. To extend the applicability of PEDRA beyond MPEs to other metastatic lesions, we generated a protocol starting from core needle biopsies. Owing to its reproducible and quantitative nature, PEDRA may provide a valuable diagnostic tool to guide time-sensitive clinical therapy decisions. Additionally, PEDRA has great potential for preclinical testing of investigational drugs, thereby reducing the need for animal experiments.

## INTRODUCTION

Precision oncology, mostly understood as therapy selection based on molecular profiling, has significantly improved outcomes for subsets of patients with actionable mutations, including NTRK fusions (*1, 2*), *ALK* translocations (*3–5*) and *EGFR* mutations (*6–10*) in NSCLC. However, the limits of this paradigm have become all too evident in recent years. Large-scale clinical programs have demonstrated variable clinical benefit of genomics-matched therapies in advanced disease (*11–16*). This is largely driven by the relative infrequency of actionable alterations, the lack of effective targeted agents for many detected drivers, and the inconsistent response observed even for well-established genomic matches (*17*). Furthermore, widely used systemic therapies such as chemotherapy and immunotherapy lack robust genomic biomarkers to guide patient selection (*18*). These challenges are exacerbated in later lines of therapy, where prior treatment exposures, clonal evolution (*19*), microenvironmental, and immune dysregulation (*20*) further diminish the predictive value of genomic profiling and complicate therapeutic decision making.

Functional precision oncology, aiming to directly measure drug responses in patient-derived tumor samples, has emerged as a complementary approach capable of revealing phenotypic sensitivities that cannot be inferred from molecular profiling alone (*21, 22*). Historically, functional drug testing relies on expanding patient tumors into cell lines or organoids. Growing evidence indicates that drug responses measured in these models may correlate with clinical outcomes in observational studies (*23–26*), yet drawbacks for their translation include their variable establishment success rates, prolonged culture times, and absence of the native stromal and immune microenvironment. More recently, rapid *ex vivo* assays that test drug responses directly in freshly obtained patient tissue have been developed, enabling instant assessment of drug efficacy and substantially reducing turnaround times for clinical decision making (*27–31*). Clinical studies demonstrate that such assays can inform treatment selection and provide meaningful benefit even in aggressive (*28, 32*) and highly refractory hematologic malignancies (*31*).

Existing functional assays are largely optimized for primary tumor specimens since they typically require substantial amounts of tissue. In advanced disease, however, only limited material is often available from metastatic lesions, such as needle biopsies, thereby restricting the feasibility and clinical integration of functional testing in this setting. Moreover, many current platforms fail to represent the autologous cellular microenvironment and instead rely on bulk readouts that can obscure context-specific drug responses. This limitation is particularly critical for therapies whose efficacy depends on cellular interactions, including immune checkpoint inhibitors (ICIs) and ADCs. Consequently, there is a need for fast yet biologically relevant and scalable platforms capable of testing multiple therapeutic modalities in metastatic tissue within clinically actionable timeframes. Here, we address this gap by developing an *ex vivo* drug-response platform to deliver functional evidence that complements molecular profiling and supports both individualized treatment selection and preclinical evaluation of emerging therapeutic modalities.

## RESULTS

### A platform to measure drug responses *ex vivo* in metastatic biopsies

To enable fast and quantitative assessment of drug responses in various metastatic patient samples, we developed a workflow that integrates automated image-based and cytokine response analyses following short-term *ex vivo* culture of patient-derived cells (Fig. 1A). Because evaluation of immunomodulatory drug effects requires preservation of autologous immune cells during *ex vivo* culture, we first optimized culture conditions using malignant pleural effusions (MPEs) from five NSCLC patients, which comprise cancer and diverse immune cell subsets (Fig. 1B). Conventional culture media (RPMI 1640, DMEM/F12) as well as the physiologically relevant human plasma–like medium (HPLM) supported overall cell viability over seven days at comparable levels (Fig. 1C). Given that HPLM recapitulates the metabolic composition of human plasma (*33*), we further evaluated this medium across all five patient-derived MPE samples. Despite inter-donor variability, HPLM maintained overall viability after five days of culture (p = 0.1383; Fig. 1D). Culturing in HPLM largely preserved therapeutically relevant cell populations, including cancer cells (p = 0.6250), T cells (p = 0.3125), and NK cells (Fig. 1E), and maintained stable expression of key therapeutic targets such as the ADC targets Nectin-4 (p = 0.6250) and TROP2 (p = 0.8750) and the immune checkpoints PD-L1 (p = 0.2500) and PD-1 (p = 0.6250; Fig. 1F). To assess whether immune cells retained functional competence, we quantified baseline cytokine secretion after five days of culture. Samples from treatment-naïve patients (P369, P206) exhibited broad cytokine secretion consistent with immune-active phenotypes, whereas samples from pretreated patients (P357, P358) showed minimal secretion (Fig. 1G, Table 1, Supplementary Table 1). These differences were independent of cellular composition (Fig. 1B), overall viability (Fig. 1D), and PD-1/PD-L1 expression (Fig. 1F).

**Table 1.** Description of treatments, clinical mutations, and outcomes of patients in the study.

| Patient ID | Sex | Age (years) | General diagnosis | Clinical mutations | PD-L1<sup>+</sup> (TPS) | Previous therapy lines | Previous treatments | Best clinical response | Following treatments | Best clinical response |
| --- | --- | --- | --- | --- | --- | --- | --- | --- | --- | --- |
| P179 | M | 39 | NSCLC | *EML4-ALK* fusion | 2% | 1 | Cisplatin + Pemetrexed + Pembrolizumab | PR | Alectinib | SD |
| P206 | M | 72 | NSCLC | *EGFR* exon 20 insertion | 5% | 0 | None | None | Cisplatin + Pemetrexed<br>Afatinib + Cetuximab | PD<br>PD |
| P357 | M | 67 | NSCLC | None | <1% | 1 | Cisplatin + Pemetrexed | SD | Docetaxel + Nintendanib<br>Nivolumab | PR<br>PD |
| P358 | F | 65 | NSCLC | *EGFR* exon 20 insertion p.H773dup | 2% | 5 | Cisplatin + Pemetrexed<br>Erlotinib<br>Docetaxel<br>Pembrolizumab<br>Osimertinib | PD<br>PD<br>PD<br>PD<br>SD | None | None |
| P369 | F | 65 | NSCLC | *EGFR* exon 19 deletion | <1% | 0 | None | None | Osimertinib | PR |

SD, stable disease; PD, progressive disease; PR, partial response

**Fig. 1.**
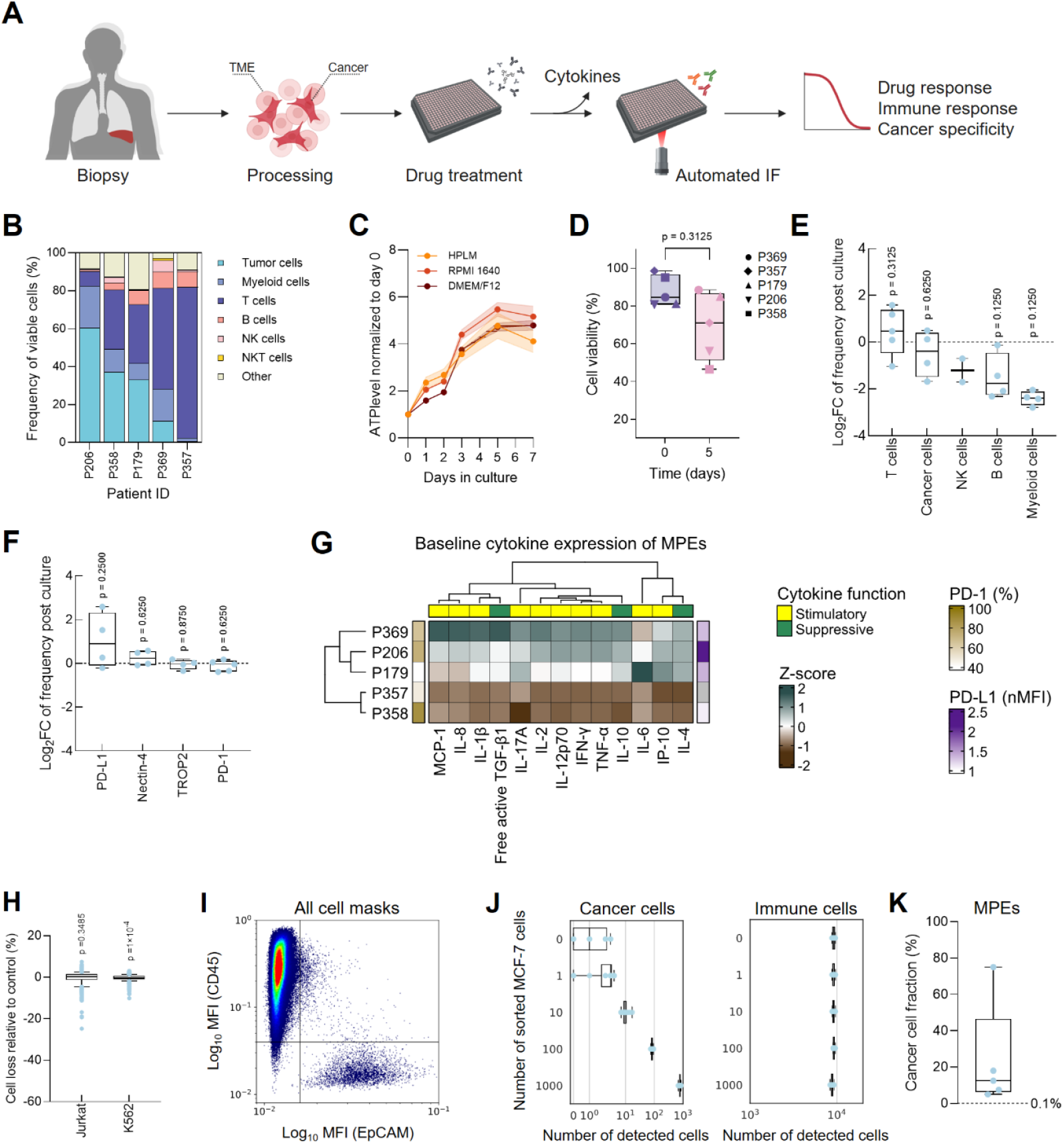
Establishment of a workflow for ex vivo drug-response profiling in metastatic biopsies. (**A**) Schematic illustration of the drug response assay. Created with BioRender.com. TME, tumor microenvironment; IF, immunofluorescence. (**B**) Cellular composition of malignant pleural effusions (MPEs) from five NSCLC patients as determined by flow cytometry. (**C**) ATP-based viability of a MPE in different culture media. Each dot represents the mean ± standard deviation (shaded envelope) of n = 5 technical replicates from one representative donor (P179). (**D**) Flow cytometric analysis of MPE viability in HPLM medium. Each dot represents an individual donor (n = 5). P-values were calculated using Wilcoxon matched-pairs signed rank test. (**E**) Log2-fold change (Log2FC) in cellular composition of MPEs after 5 days in HPLM medium compared to baseline, analyzed by flow cytometry. Each dot represents an individual donor. Samples from n = 5 donors were measured. To illustrate fold changes, cell fractions below the 2%-detection limit were excluded from analysis. P-values were determined using a one-sample Wilcoxon test against zero. (**F**). As for (E), but for marker expression of cancer cells (PD-L1, Nectin-4, TROP2) and T cells (PD-1). (**G**) Cytokine secretion and immune checkpoint expression of MPEs (n = 5) after 5 days of ex vivo culture. Color scale represents Z-scores normalized across patients for each cytokine or marker. nMFI, mean fluorescence intensity normalized to fluorescence minus one (FMO) control. (**H**) Relative cell loss of two non-adherent cell lines following the optimized IHC liquid handling protocol. Dots represent outliers among technical replicate wells across n = 3 biological replicates (384-well plates). P-values were calculated using a one-sample Wilcoxon test. (**I**) Pseudocolor plot depicting MFIs of CD45 and EpCAM signal for all segmented masks of a spike-in of MCF-7 cells in lymph node cells. Cell populations were classified based on manual gating. (**J**) Number of cancer (left) and immune cells (right) detected using our image analysis pipeline. Box plots represent data of n = 10 technical replicate wells. (**K**) Cancer cell fractions in malignant pleural effusions (MPEs) from NSCLC patients (n = 5), determined by EpCAM-based immunocytochemistry. The dashed line indicates the detection limit (10 cells per well) of the drug response assay.

For robust quantification of drug effects on both cancer cells and immune cells of clinical samples, we developed an automated immunofluorescence (IF) protocol in 384-well plates. As standard IF procedures resulted in substantial loss of non-adherent cells during liquid-handling steps (Supplementary Fig. 1A, B), we optimized the liquid handling parameters to preserve suspension cells throughout staining and imaging. Validation with non-adherent Jurkat and K562 cell lines demonstrated high cell recovery and uniform well-to-well distribution across six independent 384-well plates (Fig. 1H; Supplementary Fig. 1C, D). To reliably distinguish tumor from immune compartments at single-cell resolution, we optimized staining conditions using fluorescently labeled antibodies against EpCAM and CD45 (Supplementary Fig. 1E), enabling accurate classification of cancer and immune cells by automated image analysis (Fig. 1I).

In metastatic disease, available specimens frequently comprise samples with variable tumor cell fractions, such as MPEs (Fig. 1B), or small core biopsies that yield only minimal input material. To ensure compatibility with these specimens, we validated the sensitivity of our assay to detect rare cancer cells. Spike-in experiments mixing MCF-7 cells with patient-derived lymph node (LN) cells demonstrated reliable detection of as few as 10 cancer cells among 10,000 total cells per well (0.1%; Fig. 1J), confirming compatibility with the low tumor fractions commonly observed in MPEs, as determined by immunocytology (Fig. 1K).

### *Ex vivo* drug responses mirror observed and expected patient outcomes

To determine how well our platform recapitulates clinically observed drug sensitivities, we generated *ex vivo* drug-response profiles from all five MPEs. These cases captured a clinically relevant range of actionable genomic alterations, prior treatment exposures, and observed therapeutic outcomes (Table 1). Each MPE was screened against a 52-agent panel encompassing chemotherapies, targeted therapies, immune checkpoint inhibitors, antibody–drug conjugates, and established combination regimens of the NSCLC guideline (*34*) (Fig. 2A; Supplementary Table 2) at their maximum clinically achievable plasma concentration (*c*_max_), as defined by pharmacokinetic parameters from human trials (*35*). Assay performance met established quality standards (*36*), demonstrated by robust Z′-factors greater than 0 and signal-to-noise ratios (SNR) > 3 across patient samples (Fig. 2B). Image analysis showed high reproducibility, as determined by parallel analysis with commercially available software (MIKAIA^®^ (*37*), Supplementary Fig. 1F). In the absence of a predefined statistical cutoff requiring a large, independent validation study, drug responses were classified using empirical criteria. A compound was considered effective if it reduced cancer cell numbers relative to solvent controls (relative cancer cell reduction < 0) and exhibited preferential cytotoxicity toward cancer cells compared to immune cells, defined by a cancer-specificity score > 0 (Fig. 2C).

**Fig. 2.**
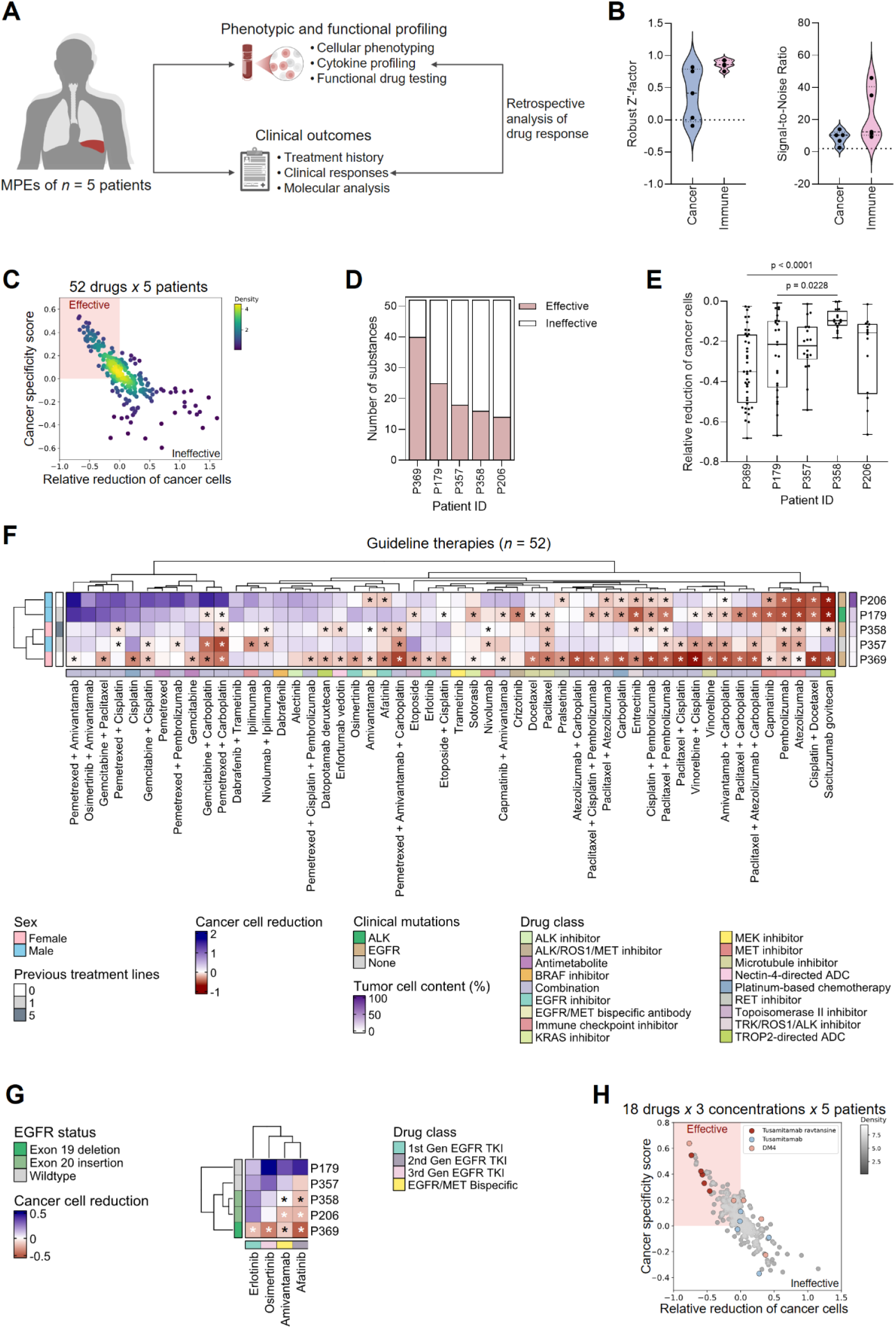
A proof-of-concept study to measure functional drug responses in MPEs. (**A**) Schematic illustration of proof-of-concept study. Created with BioRender.com. (**B**) Violin plots showing assay performance metrics for cancer and immune cell populations across MPE samples. DMSO served as negative control and benzethonium chloride as the positive control. Each point denotes one donor (n = 5). (**C**) Density scatter plot showing cancer specificity scores (y-axis) versus relative reduction of cancer cells (x-axis) of all drugs across MPEs (n = 5). Negative values of the relative reduction correspond to a reduction of cells compared to the negative control. Each dot represents one drug. Shaded quadrant indicates effective therapies. (**D**) Distribution of drug effectiveness across patients. Stacked bar plots indicate the fraction of patient samples falling into each response category. (**E**) Boxplots showing efficacy of effective substances in reducing cancer cells across patients. Each dot represents one drug. Statistical significance was assessed using the Friedman test with Dunn’s multiple comparisons post hoc test. (**F**) Guideline therapy score matrix (n = 52 drugs; columns) across patients (n = 5; rows). Scores (color scale) represent zero-centered tumor cell reduction relative to solvent control. Tumor cell content in MPEs was assessed by immunocytology. Black and white asterisks indicate cancer-specific reduction of cells (cancer specificity score >0). (**G**) As in (F), but only for EGFR inhibitors (n = 4 drugs; columns) across patient samples (n = 5; rows). *EGFR* status refers to clinical sequencing results of the respective primary tumors. Asterisks indicate cancer-specific reduction of cells (cancer specificity score >0). (**H**) Density scatter plot showing cancer specificity scores (y-axis) versus relative reduction of cancer cells (x-axis) of all drugs and concentrations of the investigational ADC panel across MPEs (n = 5). Each dot represents one drug and concentration. Quadrants indicate distinct response categories. Highlighted are tusamitamab ravtansine (50 µg/mL), its antibody control tusamitamab (50 µg/mL), and its payload control DM4 (50 µM).

MPEs demonstrated inter-patient heterogeneity in drug responsiveness, reflected in both the number of effective agents identified per sample (Fig. 2D) and the potency of effective compounds (Fig. 2E). Across the cohort, cancer cells from P369 were the most drug-sensitive, whereas cells from P358 displayed the lowest overall sensitivity (p < 0.0001; Fig. 2E). This pattern was consistent with patients’ treatment histories, as P369 was treatment-naïve and P358 had received the most extensive prior therapy at the time of MPE collection (Fig. 2F). Across all samples, sacituzumab govitecan, the cisplatin–docetaxel combination, and the immune checkpoint inhibitors atezolizumab and pembrolizumab ranked among the most active agents (Fig. 2F). We next asked whether MPEs also differed in the therapeutic classes to which they respond (Fig. 2F). We found that the treatment-naïve P369 sample was most responsive to chemotherapies while the heavily pretreated P358 showed chemoresistance. Although all MPE samples exhibited low PD-L1 tumor proportion scores (TPS <5%; Table 1), we observed sensitivity to PD-1/PD-L1–targeting immunotherapies in P357 (<1%), P179 (2%), and P206 (5%), whereas P369 (<1%) showed only minimal responsiveness. Strikingly, responses to EGFR inhibitors closely reflected the underlying *EGFR* mutation status of patients (Fig. 2G). The exon 19 variant in P369 conferred broad sensitivity to EGFR inhibition, whereas samples harboring exon 20 insertions (P206, P358) were selectively sensitive to amivantamab and afatinib, indicating that our assay may accurately capture drug class- and mutation-specific drug responses.

Given the increasing relevance of ADCs in advanced cancer treatment and the strong performance of the TROP2-targeting ADC sacituzumab govitecan in four samples, we broadened the analysis to include eight investigational ADCs (iADCs), tested alongside matched antibody-only and payload-only controls (Supplementary Table 3). This expanded ADC panel revealed further heterogeneity in drug responsiveness across MPEs (Supplementary Fig. 2).

Notably, the CEACAM5-targeting ADC tusamitamab ravtansine showed consistent and robust on-target activity in all samples, whereas the respective antibody-only control was inactive, and the free payload exhibited variable potency (Fig. 2H). Taken together, these findings provide proof-of-concept that the assay may capture both genetic and treatment-associated determinants of drug response across diverse therapeutic classes and provides initial evidence that *ex vivo* sensitivity patterns align with known clinical outcomes.

### Functional profiling reveals broad drug sensitivities in a treatment-naïve sample

To assess whether functional ex vivo testing could have provided clinically meaningful individual guidance for a treatment-naïve patient, we examined in detail the case of a patient in stage IVa with TTF-1^+^ lung adenocarcinoma (P369; Fig. 3A) and evaluated how closely the *ex vivo* sensitivity pattern aligned with the subsequent treatment course. The patient achieved a complete response to initial surgery and radiotherapy and remained in remission for 14 months before developing recurrent disease with MPE formation. Molecular profiling of the MPE identified an *EGFR* exon 19 deletion (Table 1), leading to initiation of first-line therapy with the third-generation EGFR inhibitor osimertinib, which resulted in a partial clinical response.

**Fig. 3.**
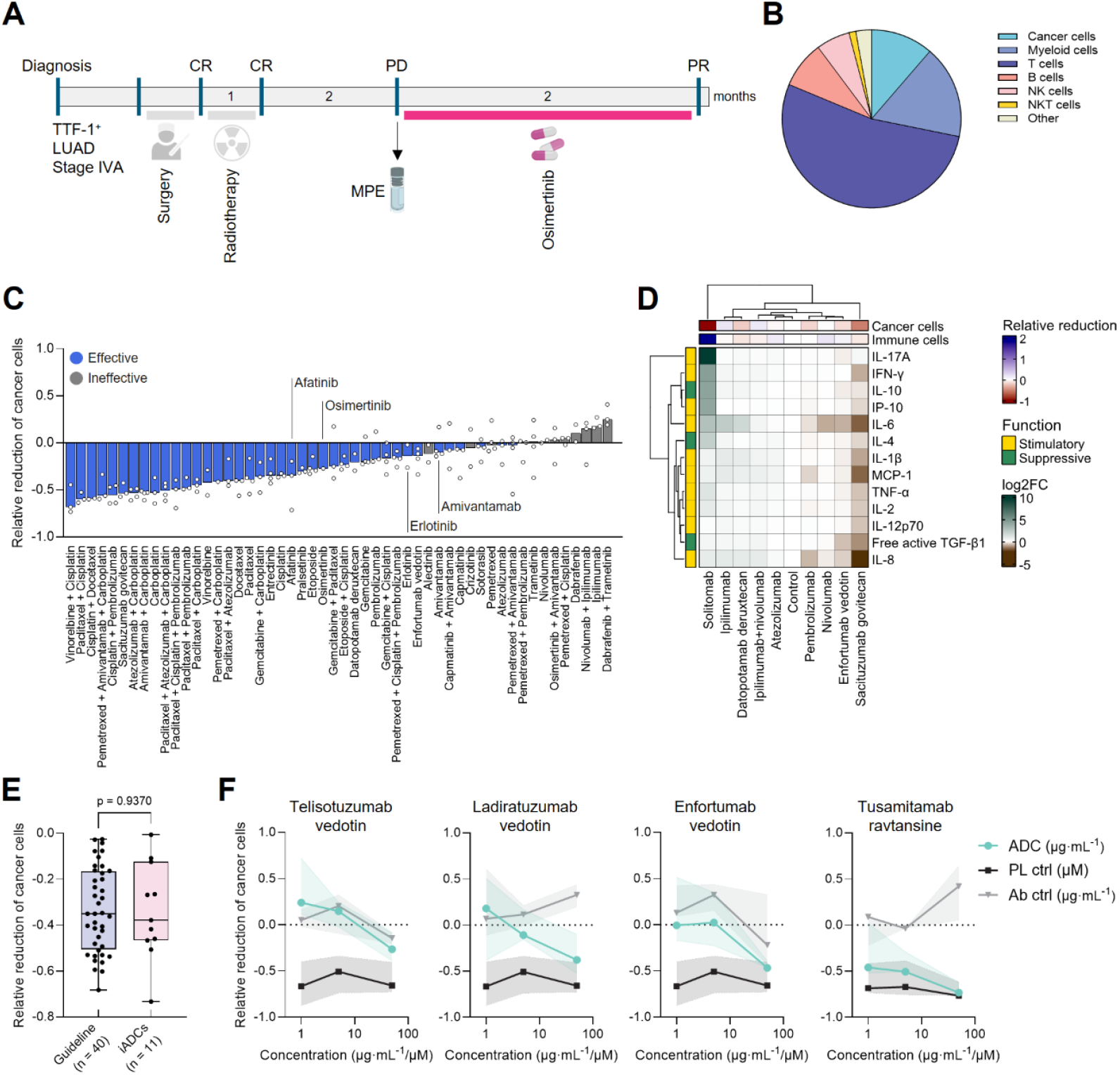
Functional drug responses in an MPE of a treatment-naïve NSCLC patient. (**A**) Clinical course of patient P369. Time intervals displayed within the bar correspond to the duration of therapy in months. CR, complete response; PD, progressive disease; PR, partial response. (**B**) Pie chart depicting immune cell composition of the MPE, based on flow cytometry analysis. (**C**) Ranking of n = 52 guideline drugs based on their ability to reduce cancer cell counts. Bars represent median. Each dot represents one technical replicate well. Blue bars indicate effective drugs; treatments clinically administered to the patient are indicated in the plot. (**D**) Cytokine secretion matrix for n = 13 cytokines (rows) across immunomodulatory drugs (n = 9; columns), measured by multiplex cytokine assay. Scores (color scale) indicate Log2-fold changes of concentrations relative to isotype control. (**E**) Box plot showing cancer cell reduction by effective drugs from the guideline and effective ADCs from the investigational ADC panel. Each dot represents one drug. Statistical significance was assessed using the Mann–Whitney test. (**F**) Dose– response curves of effective ADCs. Data are shown as median ± 95% CI of n = 2-5 replicate wells. Ab, antibody; PL, payload.

Flow cytometric phenotyping of the MPE revealed a T cell–dominant microenvironment, with T cells representing 53.1% of viable cells, followed by myeloid cells (16.9%), cancer cells (11.3%), and NK cells (8.5%; Fig. 3B). Functional drug testing identified 40 effective agents, indicating broad sensitivity to therapies of the treatment guideline. Cells exhibited pronounced sensitivity to EGFR-targeted inhibitors over other targeted therapies, aligning with the clinical choice of osimertinib and the patient’s partial response to EGFR-inhibition (Fig. 3C). Notably, the MPE also showed high sensitivity to platinum-based regimens, consistent with the treatment-naïve status of the patient. Whereas several drugs induced broad cytotoxic effects in cancer cells, immune-modulatory agents triggered only limited immune activation as determined by cytokine profiling (Fig. 3D). Only the positive-control bispecific antibody solitomab induced notable cytokine secretion, whereas the other agents produced minimal cytokine release. Although ADCs did not demonstrate higher overall potency than guideline therapies (p = 0.9040; Fig. 3E), several agents, in particular ladiratuzumab vedotin, enfortumab vedotin, and tusamitamab ravtansine, achieved effective tumor cell reduction (Fig. 3F). The cells showed marked sensitivity to ADCs linked to the microtubule-disrupting monomethyl auristatin E (MMAE) payload. Among all ADCs tested, tusamitamab ravtansine elicited the strongest antitumor activity. Thus, the assay may have supported clinical decision-making by highlighting vulnerabilities to EGFR inhibitors, platinum-based chemotherapy, and MMAE-containing ADCs, while corroborating clinical experience that patients with EGFR-mutant NSCLC derive limited benefit from ICIs.

### A refractory MPE exhibits restricted drug responsiveness ex vivo

Given the limited therapeutic options available for treatment-refractory NSCLC, we next examined whether functional *ex vivo* profiling can recapitulate a refractory state and uncover actionable residual sensitivities. One of our patient samples (P358) was obtained from a patient with stage IVa TTF-1^+^ lung adenocarcinoma after he had progressed following five prior systemic therapies, including platinum-based chemotherapy (cisplatin + pemetrexed), EGFR tyrosine kinase inhibitors (erlotinib and osimertinib), taxane-based chemotherapy (docetaxel), and PD-1 blockade (pembrolizumab; Fig. 4A). Molecular profiling conducted before and after pembrolizumab treatment confirmed an actionable *EGFR* exon 20 insertion mutation (Table 1).

**Fig. 4.**
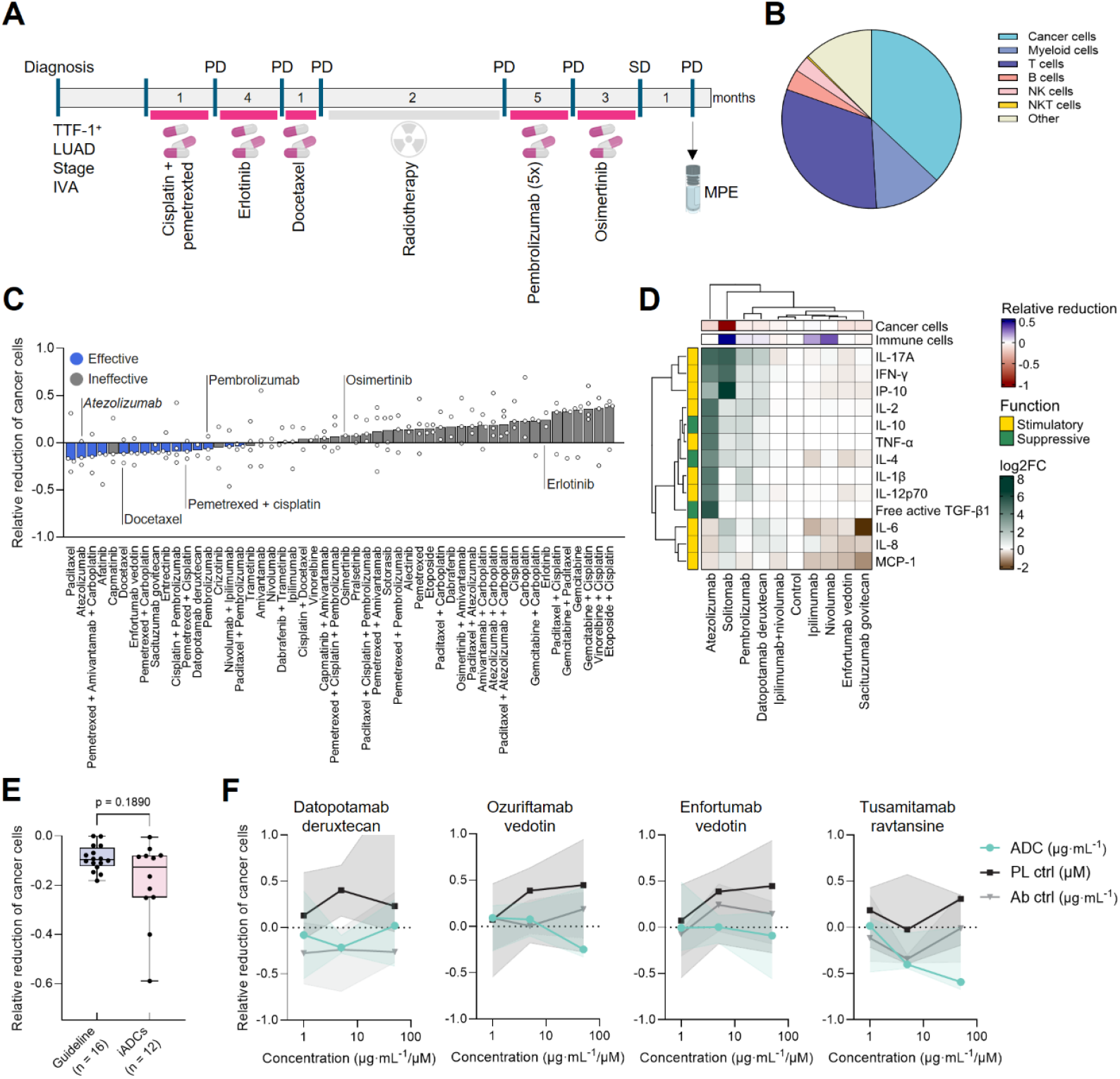
Functional drug responses in an MPE of a highly refractory NSCLC patient. (**A**) Clinical course of patient P358. Time intervals displayed within the bar correspond to the duration of therapy in months. CR, complete response; PD, progressive disease; SD, stable disease. (**B**) Pie chart depicting immune cell composition of the MPE, based on flow cytometry analysis. (**C**) Ranking of n = 52 guideline drugs based on their ability to reduce cancer cell counts. Bars represent median. Each dot represents one technical replicate well. Blue bars indicate effective drugs; treatments clinically administered to the patient are indicated in the plot. (**D**) Cytokine secretion matrix for n = 13 cytokines (rows) across immunomodulatory drugs (n = 9; columns), measured by multiplex cytokine assay. Scores (color scale) indicate Log2-fold changes of concentrations relative to isotype control. (**E**) Box plot showing cancer cell reduction by effective drugs from the guideline and effective ADCs from the investigational ADC panel. Each dot represents one drug. Statistical significance was assessed using the Mann–Whitney test. (**F**) Dose– response curves of effective ADCs. Data are shown as median ± 95% CI of n = 2-5 replicate wells. Ab, antibody; PL, payload.

Flow cytometric analysis revealed a heterogeneous immune landscape, with cancer cells comprising 37% of total events, followed by T cells (31.3%), myeloid cells (12.1%), B cells (3.8%), and NK cells (2.9%) (Fig. 4B). Notably, 87.4% of T cells expressed PD-1 (Fig. 1G). Functional drug testing identified only 16 compounds with selective cytotoxic activity, with the most active agents including paclitaxel, atezolizumab, and a triplet of pemetrexed, amivantamab, and carboplatin (Fig. 4C). However, none of the agents exhibited a relative reduction of cancer cells below −0.5, mirroring the patient’s highly refractory clinical state. This absence of activity encompassed all drugs to which the patient had previously developed resistance *in vivo*, underscoring that the *ex vivo* profile faithfully recapitulated the clinically observed treatment-refractory phenotype. Remarkably, cytokine profiling revealed a robust immunomodulatory response to atezolizumab that exceeded even the response elicited by the positive control solitomab and indicated functional engagement of the immune compartment (Fig. 4D).

Furthermore, despite resistance across multiple drug classes, a subset of iADCs demonstrated greater cancer cell reduction than the best-performing guideline-recommended therapy (Fig. 4E). Treatment with datopotamab deruxtecan, ozuriftamab vedotin, enfortumab vedotin, and tusamitamab ravtansine resulted in effective tumor cell reduction, with tusamitamab ravtansine eliciting the most pronounced effect (Fig. 4F). Notably, comparable patterns were observed in the remaining three samples, with *ex vivo* sensitivities aligning with clinical courses and mutational profiles (Supplementary Fig. 3A-C), and immunomodulatory activity—particularly to atezolizumab—emerging in an additional case (Supplementary Fig. 4A-C). Taken together, these findings show that *ex vivo* profiling can reveal residual pharmacologic vulnerabilities in refractory *EGFR*-exon 20 mutant NSCLC, particularly to ADCs, and select immune-modulatory agents.

### Feasibility of drug testing in needle biopsies of solid metastases

Since MPEs are available only from a limited subset of patients, we adapted the workflow for core needle biopsies (CNBs) of solid tumor components (e.g. metastases), which are routinely collected across late-stage malignancies. Whereas MPEs are naturally obtained as single-cell suspensions enabling accurate and scalable quantification of drug responses, needle biopsies yield small, heterogeneous tissue fragments (10–20 mg; Fig. 5A). We therefore established a protocol to homogenize and generate viable single-cell suspensions from needle biopsies, that enable precise, high-throughput drug-response profiling. Dissociation of metastatic biopsies yielded sufficient viable cells across all samples from each of six patients (median 465,000 cells; Fig. 5B), allowing for testing of multiple therapeutic candidates. Despite the limited material and required tissue digestion, IF staining yielded high-quality cellular labeling (Fig. 5C) and robust quantification of cytotoxic effects, as demonstrated by clear separation of negative and positive controls for both cancer and immune cells (p < 0.0001) with Z′-scores of 0.7 and 0.2, respectively, in a liver metastasis from a cancer of unknown primary (Fig. 5D). In this sample, testing four compounds at two concentrations was feasible, enabling reliable drug ranking, with gemcitabine and irinotecan emerging as the most potent agents (Fig. 5E). The applicability of the assay to metastasis needle biopsies was further confirmed in a bone metastasis from a breast cancer patient, where six compounds demonstrated activity and vinorelbine emerged as the most effective agent (Fig. 5E). Together, these results demonstrate that our functional drug testing workflow is compatible even with small biopsy specimens and therefore enables reliable detection of pharmacologic vulnerabilities across diverse metastatic contexts.

**Fig. 5.**
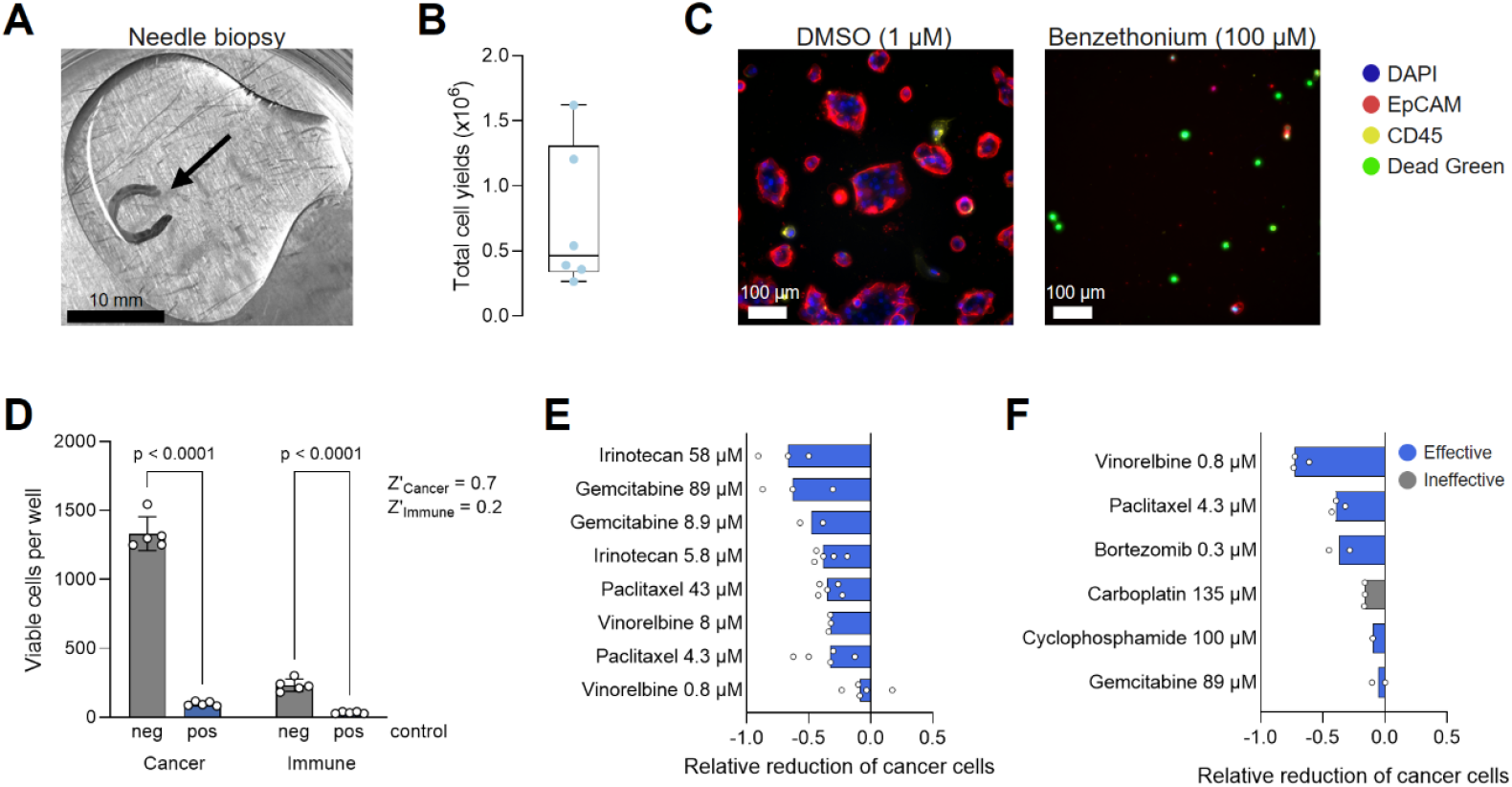
Feasibility of ex vivo drug testing in metastatic needle biopsies. (**A**) Representative image of a needle biopsy from a cancer of unknown primary (CUP) liver metastasis. Scale bar: 10 mm. (**B**) Total cell yields after homogenization of core needle biopsies (CNBs) from carcinoma patients (n = 6). Up to three CNBs were obtained per patient. (**C**) Example images of ex vivo drug response profiling of a needle biopsy of a CUP liver metastasis from negative (DMSO) and positive (benzethonium chloride) controls. Scale bar: 100 µm. (**D**) Bar graph demonstrating high assay quality in needle biopsies. Each dot represents one technical replicate well. Statistical significance was assessed using two-way ANOVA with Sidak’s multiple comparisons test. (**E**) Ranking of n = 4 drugs tested at two concentrations in a liver metastasis of a patient with cancer of CUP syndrome. Bars represent median. Each dot represents one technical replicate well. Blue bars indicate effective drugs. (**F**) Ranking of n = 6 drugs tested at single concentrations in a bone metastasis of a breast cancer patient. Bars represent median. Each dot represents one technical replicate well.

## DISCUSSION

Here, we present our patient-derived ex vivo drug response assay PEDRA that comprises high robustness, reliable quantification, and rapid generation of results. It also demonstrates high versatility to drugs of different classes, including the measurement of immune responses mediated by immunotherapies and applicability to different tissue sources. Application of the assay for different purposes, such as pre-clinical target evaluation, mode of action studies, replacement and reduction of animal experiments, and even functional precision oncology, is likely to address and overcome some significant hurdles of today’s research and clinical translation.

Functional precision oncology is beginning to influence treatment decisions in hematologic malignancies (*28, 32*) and early data now suggest that similar strategies may be applied to solid tumors, further expanding the scope of functional testing beyond hematologic cancers (*29, 30*). Historically, most *ex vivo* drug-testing efforts have depended on primary tumor specimens, largely because substantial cell numbers are traditionally required for assay performance.

However, in advanced disease, primary tumors may lose clinical relevance, as they have often been resected and metastatic cells can differ in their clonal composition due to both therapeutic pressure and endogenous selection mechanisms and ultimately drive disease progression. Access to metastatic material suitable for functional drug testing remains challenging.

MPEs can provide comparatively large amounts of viable material in a subset of patients with advanced solid malignancies (∼15%) (*38, 39*), but their cellular composition and tumor cell content varies substantially between patients (*30, 40–42*). In contrast, needle biopsies of metastatic lesions are rarely available beyond immediate histopathological or molecular analyses because they yield only minimal tissue and are characterized by pronounced heterogeneity within individual samples. To address these challenges, we established a highly robust workflow that is compatible with both heterogeneous cell mixtures and low total cell numbers. Dissociation of core needle biopsies (CNBs) enabled parallel testing of multiple agents from limited starting material, whereas microenvironment-preserving methods such as tissue slice cultures are often constrained by substantial tissue requirements and the risk of sampling bias arising from intratumoral heterogeneity (*43*). Because CNBs represent a routine diagnostic procedure for evaluating metastatic disease across virtually all cancer types (*44*), this compatibility substantially broadens the platform’s clinical applicability and demonstrates that functional drug testing could possibly be integrated into routine diagnostic workflows of advanced cancer patients in the future.

The therapeutic landscape of metastatic cancer is increasingly dominated by next-generation agents such as antibody–drug conjugates and immunotherapies. Because their activity critically depends on the tumor microenvironment, particularly on the presence and function of immune cells, *ex vivo* assays must preserve the phenotypic integrity of all relevant cellular compartments, allow sufficient incubation time to capture delayed immune-mediated effects, and incorporate readouts capable of quantifying these responses with, sensitivity, specificity, and robustness. Our assay preserved the native cellular composition of metastatic samples for a duration sufficient to capture both cytotoxic and immunomodulatory drug effects (*45, 46*), and supported high-throughput testing within a timeframe (1-2 weeks) compatible with clinical decision-making.

This is in stark contrast to approaches that expand selected cell populations (*47, 48*) or rely on coculture systems to model the microenvironment (*49, 50*). By integrating cell lineage-resolving imaging with multiplex cytokine profiling, our platform enables evaluation of the efficacy of all major drug classes within the same patient sample.

Of further interest is the adequate representation of clinical disease courses over therapy *in vitro*. Platinum-doublet chemotherapy remains the standard first-line treatment for NSCLC tumors without actionable mutations (*34, 51*). Yet these regimens are still administered without reliable biomarkers. Here, we provide evidence that our assay can capture clinically relevant patterns of chemotherapy sensitivity and resistance in MPEs. We found that the clinical pattern of chemotherapy resistance in our cohort was mirrored *ex vivo* by marked variability in responsiveness to chemotherapies and platinum-based regimens, consistent with whether patients were treatment-naïve or had previously developed refractory disease. These findings suggest that *ex vivo* profiling of MPEs may offer a practical means to anticipate chemotherapy responses and support more informed therapeutic decision-making in advanced NSCLC.

In addition, mutation-based therapy escape (*52*) could be modelled by our assay. Selecting the most effective inhibitor among multiple drug generations or identifying the most responsive alteration among several targetable mutations therefore represents an important opportunity for functional assays to support precision oncology. PEDRA not only captured genotype-specific therapeutic vulnerabilities in MPEs but also discriminated between variants arising at distinct mutational loci. Consistent with clinical observations, we recapitulated the well-established sensitivity of EGFR exon 19 deletions (*53*) and the resistance of EGFR exon 20 insertions to EGFR TKIs (*54*). Importantly, we identified the EGFR–MET bispecific antibody amivantamab as a potential therapeutic option in exon 20–insertion cases, mirroring its clinical indication (*10*) and providing initial evidence that the assay can reveal actionable alternatives for TKI-resistant variants.

Beyond chemotherapies and targeted agents, the data also support the ability of our platform to detect responses to immunomodulatory therapies. The assay mirrored clinical resistance to PD-1 blockade, showing minimal *ex vivo* response in non-responding patients. Notably, the PD-L1– targeting antibody atezolizumab induced robust cytokine secretion in samples refractory to PD-1 inhibitors, suggesting that PD-L1 blockade may partially bypass mechanisms of PD-1 resistance. Although clinical evidence supporting PD-L1 therapy after PD-1 progression remains limited, these results are consistent with reports highlighting immunomodulatory effects of PD-L1 inhibition on myeloid populations (*55*) and point toward the possibility that PD-L1 blockade may provide additional benefit in selected cases.

Finally, given the increasing clinical relevance of ADCs in NSCLC—particularly in HER2-positive (*56, 57*) and EGFR-mutant (*58*) disease—we evaluated eight clinically approved or investigational ADCs. Among these, sacituzumab govitecan (TROP2-targeted) and tusamitamab ravtansine (CEACAM5-targeted) demonstrated consistent *ex vivo* activity across patient samples. These results are congruent with clinical benefit observed with sacituzumab govitecan in the phase III EVOKE-01 study (*59*) and the promising outcomes from phase I/II trials of tusamitamab ravtansine in heavily pretreated NSCLC with high CEACAM5 expression (*60*).

Thus, our assay may capture the therapeutic potential of investigational ADCs and help nominate treatment strategies for highly refractory patients. Together, these findings demonstrate that *ex vivo* drug testing of metastatic biopsies can reveal individualized vulnerabilities across multiple therapeutic classes, offering a versatile framework to advance functional precision oncology.

Of note, this study has several limitations. First, the cohort size was very small and larger, prospectively acquired datasets will be required to rigorously evaluate the generalizability and predictive performance of our assay across diverse cancer types, genomic contexts, prior treatment exposures, and clinical trajectories. Second, while tissue dissociation is required for robust measurement of drug responses and mitigates tissue sampling error in solid biopsies, this approach necessarily results in loss of spatial context, which should be considered when testing drugs whose activity depends on spatial, metabolic, or vascular features of the tumor microenvironment. Third, while the assay’s rapid turnaround of 1-2 weeks enables timely functional drug testing, insight into clonal adaptation and evolutionary dynamics that arise under sustained therapeutic pressure requires longitudinally repeated application of the assay. Finally, although the assay enables detection of immunomodulatory activity, it does not recapitulate key *in vivo* features of system-wide immune engagement, including antigen spread, T-cell recruitment, and myeloid remodeling. Combining this platform with immune-relevant tissues, such as lymph nodes, may help delineate which aspects of immune contexture can be inferred from such integrated approaches.

Future work should evaluate the assay in larger, prospective, controlled, longitudinal clinical trials embedded in a molecular tumor board and may additionally incorporate transcriptomic, (phospho-)proteomic, and spatial analyses to more comprehensively dissect mechanisms of response and resistance. Together, this information may translate into novel ways of therapy selection in a truly personalized way.

## MATERIALS AND METHODS

### Study design

The objective of this study was to establish and validate an ex vivo drug testing platform, enabling drug efficacy testing in metastatic biopsies. Therefore, we collected malignant pleural effusions (MPEs) of n = 5 NSCLC patients and core needle biopsies (CNBs) of n = 6 patients of different entities. All study protocols were approved by the Ethics Committee of the University of Regensburg (reference numbers 17-672 and 18-948-101) and adhered to the principles outlined in the current version of the Declaration of Helsinki (DoH). Written informed consent was obtained from all participating patients. Samples were collected as part of routine diagnostic or therapeutic procedures at the University Hospital Regensburg and the Clinical Center Passau. Tumor stage and grading were determined according to the international system for the staging of cancer. Data from experiments were excluded in cases of technical errors that rendered them invalid. The number of biological and technical replicates (n) is specified in each figure legend. All data points represent independent observations or donors, with no repeated measures included. If not explicitly stated, the experiment was conducted once with the presented number of data points. Plate maps of drug screening assays were fully randomized to exclude plate patterns and edge effects.

### Collection and purification of cells from metastatic samples

Pleural effusions were drained from lung cancer patients during thoracentesis or tumor resection. To isolate the cellular fraction, the effusions were centrifuged at 500 × g for 10 min at room temperature, the pellet was washed with HBSS, and cells were either enriched for mononuclear cells by density gradient centrifugation or used without enrichment. Red blood cells were removed by ACK lysis (Thermo Fisher Scientific, #A1049201) if they represented >40% of the cells. Cells were cryopreserved and stored in liquid nitrogen until usage.

Needle biopsies obtained from distant metastases under ultrasound guidance were placed in a cell culture dish, submerged in tumor digestion medium (10 mM HEPES (Sigma Aldrich, #H0887), 10,000 U/mL Penicillin, 10 mg/mL Streptomycin (Pan Biotech, #P06-07100), 1% (w/v) BSA, 0.33 mg/mL Collagenase (Sigma Aldrich, #C9407), 100 µg/mL Hyaluronidase (Sigma Aldrich, #H4272), 0.1 mg/mL DNAse I (Roche, #11284932001) in DMEM/F12 (Pan Biotech, #P04-41500) and minced into fragments of approximately 1–3 mm^3^ using sterile scalpels. Upon incubation at 37 °C for 20–40 minutes, cells were pelleted by centrifugation at 300 × g for 10 minutes at RT and dissociated in prewarmed (37 °C) trypsin/EDTA for 5 min at RT. After neutralization, the cell suspension was passed through a 40-µm cell strainer with residual tissue triturated using a syringe plunger to maximize cell recovery. Cells were centrifuged at 300 × g for 5 min at RT and cells were freshly used for drug screening assays. For both pleural effusions and needle biopsies, viable cells were enriched using magnetic beads (Miltenyi Biotec, #130-090-101) if viability prior to downstream assays was < 80%.

### Flow cytometry

Cells of MPEs were resuspended in blocking buffer (1:100 TruStain FcX™ (BioLegend, #422302) in DPBS), incubated for 10 min at 4°C, and centrifuged at 1,000 × g for 2 minutes at 4 °C. Cells were resuspended in 100 µL of FACS buffer (1% (w/v) BSA, 2 mM EDTA in DPBS) per 1 x 10^6^ cells and stained using the following antibodies: Alexa Fluor® 488 anti-human CD3, clone HIT3a (BioLegend, #300319, RRID:AB_493690), PerCP/Cyanine5.5 anti-human CD14, clone HCD14 (BioLegend, # 325621, RRID:AB_893252), PE anti-human CD279 (PD-1), clone EH12.2H7 (BioLegend, # 329905, RRID:AB_940481), PE/Cyanine7anti-human CD11c, clone Bu15 (BioLegend, # 337215, RRID:AB_2129791), APC-Vio770 anti-human CD19, clone LT19 (1:50; Miltenyi Biotec, # 130-098-073, RRID:AB_2661296), Brilliant Violet™ 421 anti-human CD45, clone 2D1 (BioLegend, # 368522, RRID:AB_2687375), Brilliant Violet™ 605 anti-human HLA-DR, clone L243 (BioLegend, # 307639, RRID:AB_11219187), Brilliant Violet™ 785 anti-human CD56 (NCAM), clone 167 (BioLegend, # 362549, RRID:AB_2566058), Alexa Fluor® 488 anti-human CD326 (EpCAM), clone Co17-1A (BioLegend, # 369808, RRID:AB_2650905), PE anti-human TROP2, REAfinity™ (1:50; Miltenyi Biotec, # 130-115-097, RRID:AB_2726914), APC anti-human Nectin-4, REAfinity™ (1:50; Miltenyi Biotec, # 130-116-103, RRID:AB_2727350), Brilliant Violet™ 421 anti-human CD274 (B7-H1, PD-L1), clone 29E.2A3 (BioLegend, # 329714, RRID:AB_2563852), Brilliant Violet™ 785 anti-human CD45, clone HI30 (BioLegend, # 304048, RRID:AB_2563129), Zombie Aqua™ (1:1,000; BioLegend, # 423102), Zombie NIR™ (1:500; BioLegend, # 423106). Unless otherwise stated, all antibodies were diluted 1:100. Samples were acquired on a CytoFLEX SRT Benchtop Cell Sorter (Beckman Coulter). Data were analyzed in FlowJo™ (BD Biosciences). Gating excluded debris (FSC/SSC), doublets (FSC-H vs FSC-A), and dead cells (via viability dye). Cell populations were defined based on marker positivity.

### Assessment of cell loss during liquid handling

To quantify cell loss attributable to automated liquid handling, Jurkat or K562 cells were resuspended at 2 x 10^5^ cells/mL in culture medium supplemented with 5 µg/mL Hoechst 33342 (Sigma-Aldrich, #14533) and dispensed at 50 µL per well (1 x 10^4^ cells per well) into 384-well plates (Greiner Bio-One #781866) using a Multidrop Combi dispenser (Thermo Fisher Scientific). Cells were allowed to settle for 45 min at RT prior to further processing. To mimic the IF procedure, 15 µL of 4% paraformaldehyde (PFA) in DPBS was added directly to each well and incubated for 15 min at RT. The fixative-containing medium was removed, 50 µL DPBS was added (blocking step), aspirated, and replaced with 20 µL DPBS (staining step). Cells were then washed twice with 50 µL DPBS. Plates were imaged by high-content microscopy immediately before fixation and after completion of the workflow. Nuclei were quantified by automated image analysis using Harmony^®^ software (Revvity; RRID:SCR_023543), and cell recovery was calculated to determine cell loss associated with the liquid handling procedure.

### Assay sensitivity analysis

Lymph nodes (LNs) from melanoma patients were obtained, processed, and screened for disseminated cancer cells as previously described (*61*). LN-cells were resuspended at 2 x 10^5^ cells/mL in culture medium and dispensed at 50 µL per well (1 x 10^4^ cells per well) into Greiner Screenstar 384-well plates using a Multidrop Combi dispenser. Cells were allowed to settle for 45 min at RT. Subsequently, 1, 10, 100, or 1,000 MCF-7 cells were sorted into each well on top of the LN cell suspension using a CytoFLEX SRT benchtop cell sorter. After an additional 60 min settling at RT, plates were processed by automated immunocytochemistry, followed by high-content microscopy and image analysis.

### Preparation of library plates

Standardized compound library plates were prepared using a D300e Digital Dispenser (Tecan Group) according to the manufacturer’s recommendations in 384-well plates. The guideline library consisted of n = 52 drugs or drug-drug combinations, respectively, tested in technical quadruplicates. The ADC library included n = 8 ADCs, n = 8 antibody controls, and n = 3 payload controls in technical quintuplicates, tested in three-point serial dilutions including 1, 5, and 50 µg/mL (ADCs and antibodies) and 1, 5, and 50 µM (payloads). All plates were prepared following a randomized plate layout to minimize plate effects with at least 10% of the wells designated as solvent (negative) controls. Benzethonium chloride (100 µM) was included as positive control. Solitomab was used as a positive control for immunomodulatory effects. Prior to drug screening, pre-spotted library plates were thawed for 30 min in the dark, and 30 µL of medium was added per well using a Multidrop Combi dispenser. Water-soluble compounds, antibodies, and ADCs were diluted to 3 mg/mL in DPBS, supplemented with 0.3% Tween-20, and freshly dispensed.

### Short-term culture and ex vivo drug treatment

Pleural effusions were thawed, dead cells removed, and viable cells were resuspended in Human Plasma-Like Medium (Thermo Fisher Scientific #A4899102) supplemented with 10% human AB serum (BioIVT #HUMANABSRMP-1), 1% Penicillin/Streptomycin (Pan Biotech #P06-07100), and 60.8 µM 2-hydroxybutyric acid (Thermo Fisher Scientific #A18636.14), and seeded in 30 µL at a density of 5 x 10^4^ cells per well. If pleural effusions contained > 80% lymphocytes, cells were seeded at 1 x 10^5^ cells per well. Needle biopsies were digested, resuspended in culture medium, seeded at 5 x 10^4^ cells per well in 50 µL, and drugs were freshly dispensed on top of the cells. Plates were sealed with gas-permeable sealing foil (Greiner Bio-One, #676050) and incubated at 37 °C in a 5% CO_2_ humidified atmosphere for 5 days.

### Automated immunocytochemistry

Following drug incubation, 50 nM Image-IT DEAD Green viability dye was added to each well and plates were incubated at 37 °C for 30 min. Cells were fixed by adding 15 µL of 4% PFA in DPBS directly to the wells, followed by a 15 min incubation at RT. After fixation, the fixative-containing medium was removed and non-specific binding was blocked with 50 µL blocking buffer (Human TruStain FcX™, BioLegend #422302; 1:200 in DPBS) for 60 min at RT. The blocking solution was aspirated, and cells were stained overnight at 4 °C with a cocktail of antibodies in 20 µL per well: APC anti-human CD326 (EpCAM), REAfinity™ (1:200; Miltenyi Biotec, Cat# 130-111-000, RRID:AB_2657497), PE/Dazzle™ 594 anti-human CD45, clone HI30 (1:200; BioLegend, Cat# 304052, RRID:AB_2563568), and DAPI (10 µM; Sigma-Aldrich #D9564), prepared in blocking buffer. After staining, cells were washed twice with 50 µL of DPBS and stored in 50 µL of DPBS at 4 °C until imaging.

### High-content microscopy

Automated fluorescence microscopy was performed using an Operetta^®^ CLS High-Content Analysis System (Revvity) or an Operetta^®^ High-Content Analysis System (Revvity), respectively. Cells were imaged using a 20x 0.4 NA air objective, capturing 4×4 or 5×5 non-overlapping fields so that more than 90% of the well surface area was covered. The number of measured z-stacks was adapted individually for each sample and measured in 5 µm distances. The images were sequentially taken from the Brightfield, DAPI (nuclei), GFP (DEAD Green, non-viability/cell death), Alexa Fluor 594 (CD45, immune), and APC (EpCAM, cancer) channels, with lasers and bandpass filter sets set so that the channels were non-overlapping.

### Image preprocessing

The raw .tiff images were transferred from the microscope and analyzed using Cellprofiler 4.2.8 with installed plugins (*62*). The z-stack images of fluorescent channels were merged using maximum projection algorithm (minimum projection for brightfield images). The resulting projection images were exported as .tiff files for further analysis. For each individual single-channel image, illumination correction was performed using a Gaussian or polynomial function. Given that each well was imaged in its entirety (including regions outside the images), the areas inside the wells were determined by thresholding the brightfield signal.

### Cell segmentation and classification

Pre-processed images of DAPI (nuclei), GFP (non-viability/cell death), Alexa Fluor 594 (immune), and APC (cancer) channels were segmented using the Cellpose cyto2 model (*63*). To segment cells of different sizes, morphologies, and growth patterns (e.g. adherent cells, clusters), threshold- and deep-learning-based algorithms were combined. Small and large single cells were detected by Cellpose cyto2 (*63*) at two separate diameter settings. The precise parameters of the Cellpose model (flow_threshold, cellprob_threshold) were adapted on a per-sample basis to account for variations in intensity, morphology, and cell growth. Adherent cells were detected using threshold-based algorithms. Additionally, the DAPI channel was used as seeds along with propagate function to segment clustered and clumped cells accurately. Standard CellProfiler intensity, texture, shape and morpholgy features were measured from nuclei, immune, and cancer cells, where applicable, and segmentation masks were exported to .tiff files. For quick visual evaluation, all images of each plate were rescaled, merged, and colorized per field and per well, and single cell crops for immune and cancer cells were generated based on their respective segmentation masks.

To classify cells, all object features and their respective neighbors were exported to SQLite files and subsequently analyzed using custom R scripts. The exported object features were merged with the relevant metadata and normalized to the corresponding solvent control values. Cancer and immune cells exhibiting either very low object intensities or very small areas were excluded based on manual gating as these cells likely presented segmentation artifacts or debris.

Additionally, outlier wells exhibiting abnormally high or low intensities or cell numbers were removed if the observed pattern could be attributed to technical errors, such as liquid handling artifacts. Marker positivity and viability of cells were determined based on manual gating per plate using mean fluorescence intensities or dimensionality reduction (UMAP) with k-means clustering of mean cell/cell membrane fluorescence intensities. Classification efficiency was curated based on negative and positive controls.

### Determination of drug responses

The relative reduction of viable immune and cancer cells per well compared to negative controls was determined based on formula (1), wherein a negative value of the relative reduction indicates a reduction of cells compared to the negative control.

1. Relative cell reduction = (number of viable cells in drug-treated well/median number of viable cancer cells in negative control wells) – 1 Cancer-specific effects of drugs were determined using formulas (2) and (3), wherein a positive cancer specificity indicates a specific decrease in cancer cell fraction compared to the immune cell fraction and negative control.
2. Cancer cell fraction = (number of viable cancer cells per well)/(number of viable cancer cells per well + number of viable immune cells per well))
3. Cancer specificity score = 1 – (cancer cell fraction in drug-treated well/median cancer cell fraction in negative control wells)

### Cytokine analysis

Fifteen µL of culture media per well were collected from assay plates after 5 days of drug incubation and immediately frozen at −80 °C. Cytokine analysis was performed using the LEGENDplex™ Human Essential Immune Response Panel (13-plex) kit in a V-bottom plate format (BioLegend # 740930), following the manufacturer’s protocol. Fluorescence data were acquired using a Gallios Flow Cytometer. Bead populations were identified based on size and internal fluorescence, and a minimum of 300 beads per analyte were acquired. Cytokine concentrations were calculated using the LEGENDplex™ Data Analysis Software.

### Statistical analysis

Details of specific statistical tests employed are provided in the respective figure legends. Data sets were assessed for normality (Gaussian distribution) between groups prior to statistical testing by Shapiro-Wilk-Test. All statistical analyses and data visualizations were performed using GraphPad Prism, R software, or Python package seaborn.

## List of Supplementary Materials

Materials and Methods Fig. S1 to S4

Table S1 to S3 Reference (*64*)

## Acknowledgments

We thank the patients and their families for making this study possible. We thank Theresa Suckert, Sabine Tyrra, Anthea Povall, Justin Skotnitzki, Nathalie Drexler, Judith Proske, and Silvia Materna-Reichelt for their assistance and support.

## Funding

Deutsche Forschungsgemeinschaft (DFG, German Research Foundation) grant SFB-TRR 305-B13 (CW), Z02 (CAK)

Bavarian Cancer Research Center (BZKF) grant CUP-PEDRA (BGF/24/12/A/KUBU) Bavarian Cancer Research Center (BZKF) grant ADC (RC)

Bavarian Cancer Research Center (BZKF) grant LT-OMICS (RC)

Bavarian Ministry of Economic Affairs, Energy and Technology (AZ 11-27999) (CAK)

Federal Ministry of Research, Technology and Space (BMFTR), GO-Bio initial grant 16LW0611 (LW)

Fraunhofer Cluster of Excellence for Immune-Mediated Diseases CIMD grant PEDRA 2.0 (CW)

## Author contributions

Conceptualization: LW, CW, AS, CK

Data curation: LW, DD, ST, KW, FL, MH, DH

Formal analysis: LW, DD, MH, VB, MB

Funding acquisition: CAK, CW, LW, BK

Investigation: LW, CW, CB, DD, CEB, DAH

Methodology: LW, CB, DH, AS, DD, MH, MB, VB, KW, NSG

Project administration: CW, LW, CAK

Resources: FL, JL, TN, TS, BK, RC, TB, MWK, NF, CAK

Software: DD, MH, MB, VB

Supervision: CW, CAK

Validation: LW, CB, MH, VB, MB

Visualization: LW

Writing – original draft: LW

Writing – review & editing: LW, DD, KW, ST, CB, DAH, AS, CEB, MH, NSG, FL,

DH, JL, TN, TS, BK, RC, MB, VB, CAK, CW

