## Supplementary Materials for "Ex vivo drug testing in metastatic biopsies reveals patient-specific vulnerabilities to cancer targeting and immune activating drugs"

#### MATERIALS AND METHODS

##### Cell culture

Jurkat, K562, and MCF-7 cells were maintained in RPMI 1640 supplemented with 10% FBS (Pan Biotech #P40-37500) and 1% Penicillin/Streptomycin and cultured at 37 °C in a humidified incubator with 5% $CO_2$.

##### Cell viability assay

Pleural effusions were thawed, dead cells removed, and viable cells were resuspended in Human Plasma-Like Medium (Thermo Fisher Scientific #A4899102), RPMI 1640 (Pan Biotech #P04-17500), or DMEM/F12 (Pan Biotech #P04-02500) supplemented with 10% human AB serum (BioIVT #HUMANABSRMP-1) and 1% Penicillin/Streptomycin (Pan Biotech #P06-07100) in 40 µL at a density of $5 \times 10^4$ cells per well ) into Greiner Screenstar 384-well plates (Greiner Bio-One #781866). HPLM-based media were supplemented with 60.8 µM 2-hydroxybutyric acid (Thermo Fisher Scientific #A18636.14). Cellular viability after culture was assessed using an ATPlite 1step Luminescence Assay System (Revvity #6016731) following the manufacturer's protocol. Thirty µL of ATPlite solution was added to each well using a Multidrop Combi Dispenser (Thermo Fisher Scientific). After vigorous mixing on an orbital shaker for 2 min, luminescence was measured using an EnVision Xcite 2105 multimode plate reader (Revvity).

##### Cell segmentation and classification with MIKAIA®

Cell segmentation and classification were performed on preprocessed images using the FL Cell Analysis App in MIKAIA® (Fraunhofer IIS, version 2.4.1, (*37*)). Segmentation was based on the DAPI (nuclei), Alexa Fluor 594 (immune) and APC (cancer) channels. Prior to analysis, each channel underwent linear contrast stretching for each multi well plate, the per channel minimum was set to the average 0.1st percentile across all wells and the maximum to the average 99.9th percentile; intensities were then linearly rescaled and clipped to [0,1]. The DAPI channel and a composite of the Alexa Fluor 594 and APC channels were subsequently provided to the AI model using MIKAIA's "Membrane + Nuclear Marker XXL AI" mode. The underlying architecture of the AI model is Cellpose-SAM (*64*) and was trained on a proprietary dataset using SAM default parameter as initialization. For each cell, both nuclear and whole cell boundary annotations were generated.

Cell classification was based on per cell marker expression within specified compartments. MIKAIA allows selection of the compartment to be evaluated, and the fraction of pixels included in the intensity average. Pixels are ranked by intensity, and the chosen fraction is averaged to yield the marker expression. EpCAM was evaluated over the whole cell area using the top 50% of pixels, whereas DEAD Green and CD45 were evaluated within the nuclear compartment using the top 70% of pixels. Cells were labeled positive for a given marker if the average expression exceeded a predefined threshold. Uniform thresholds were applied across wells within each plate and were determined from a subset of wells per plate. Final cell types were assigned automatically based on marker co expression.

#### SUPPLEMENTARY FIGURES

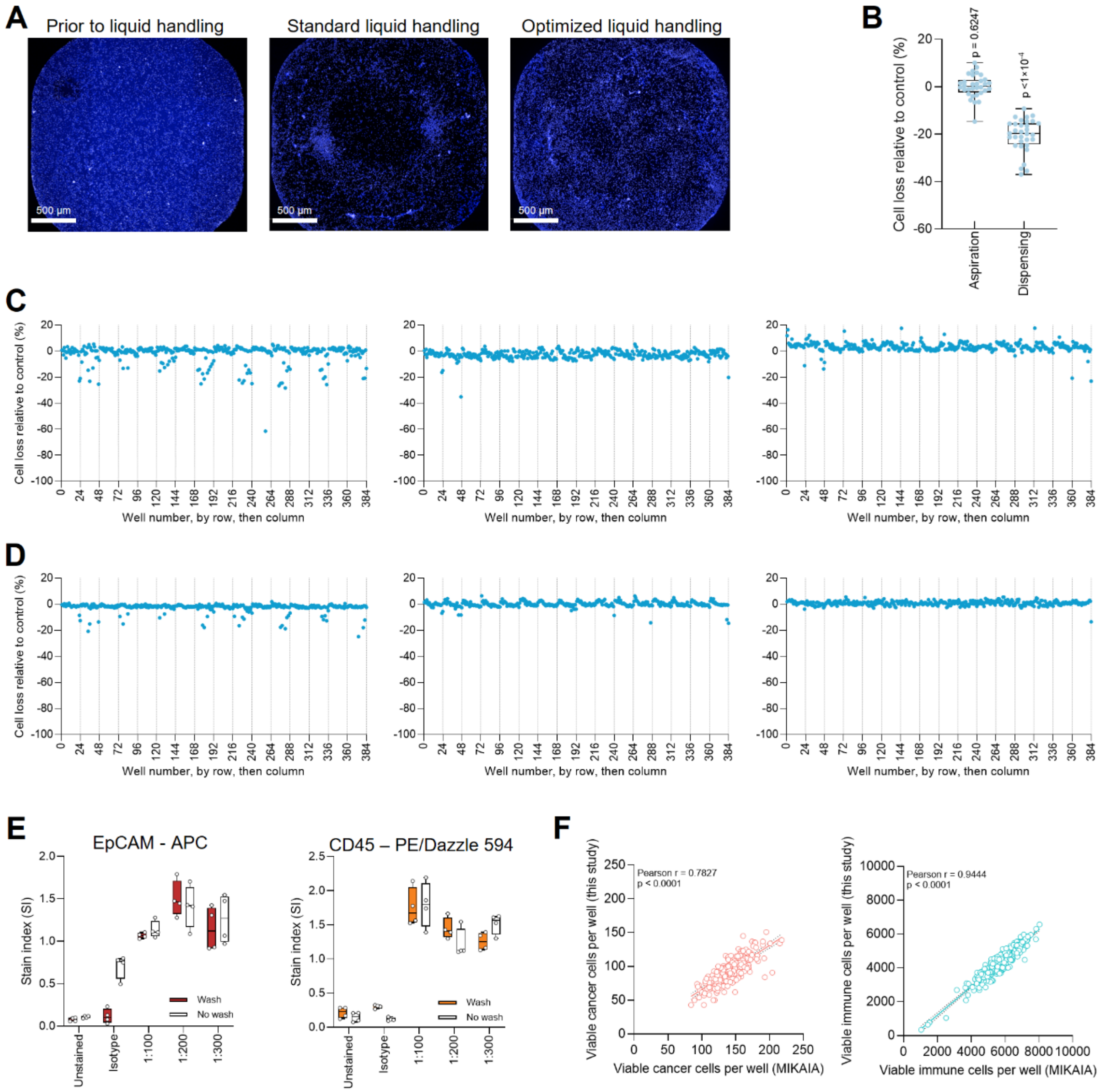


**Supplementary Fig. 1. Optimization of immunohistochemistry of non-adherent cells in 384-well plates.** (**A**) Representative whole-well IF images of PBMCs before and after standard and optimized liquid handling procedures for IF. Blue color indicates nuclear staining. Scale bar: 500 µm. (**B**) Quantification of relative cell loss (PBMCs) after liquid aspiration and dispensing. Each dot represents number of nuclei per technical replicate well (n = 32). P-values were determined using a one-sample Wilcoxon test against zero. (**C**) Cell loss (Jurkat) relative to control plotted for each well (dot) of a 384-well plate. Each graph represents one replicate 384-well plate (n = 3). (**D**) As for (C), but for K562 cells. (**E**) Stain index of MCF-7 cells labeled with varying concentrations of EpCAM-APC (left) and PBMCs stained with CD45-PE/Dazzle 594 (right), comparing washed and no-wash conditions. A dilution of 1:200 and additional washing steps were applied to both antibodies. Dots indicate technical replicate wells (n = 4). (**F**) Correlation between drug efficacy quantified with the present image-analysis workflow (y-axis) and the reference software MIKAIA (x-axis). Each dot represents a single well of the MPE of P357 treated with the ADC drug panel.

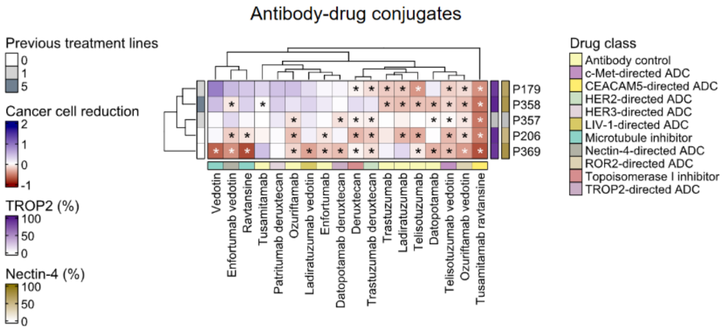


**Supplementary Fig. 2. Measurement of MPE responses to antibody-drug conjugates.** Antibody-drug conjugate score matrix (n = 7 antibody-drug conjugates, n = 7 antibody controls, n = 3 payload controls; columns) across patients (n = 5; rows). Scores (color scale) represent zero-centered tumor cell reduction relative to solvent control. Target expression was quantified by flow cytometry. Asterisks indicate cancer-specific reduction of cells (cancer specificity score >0). Only highest concentrations are shown.

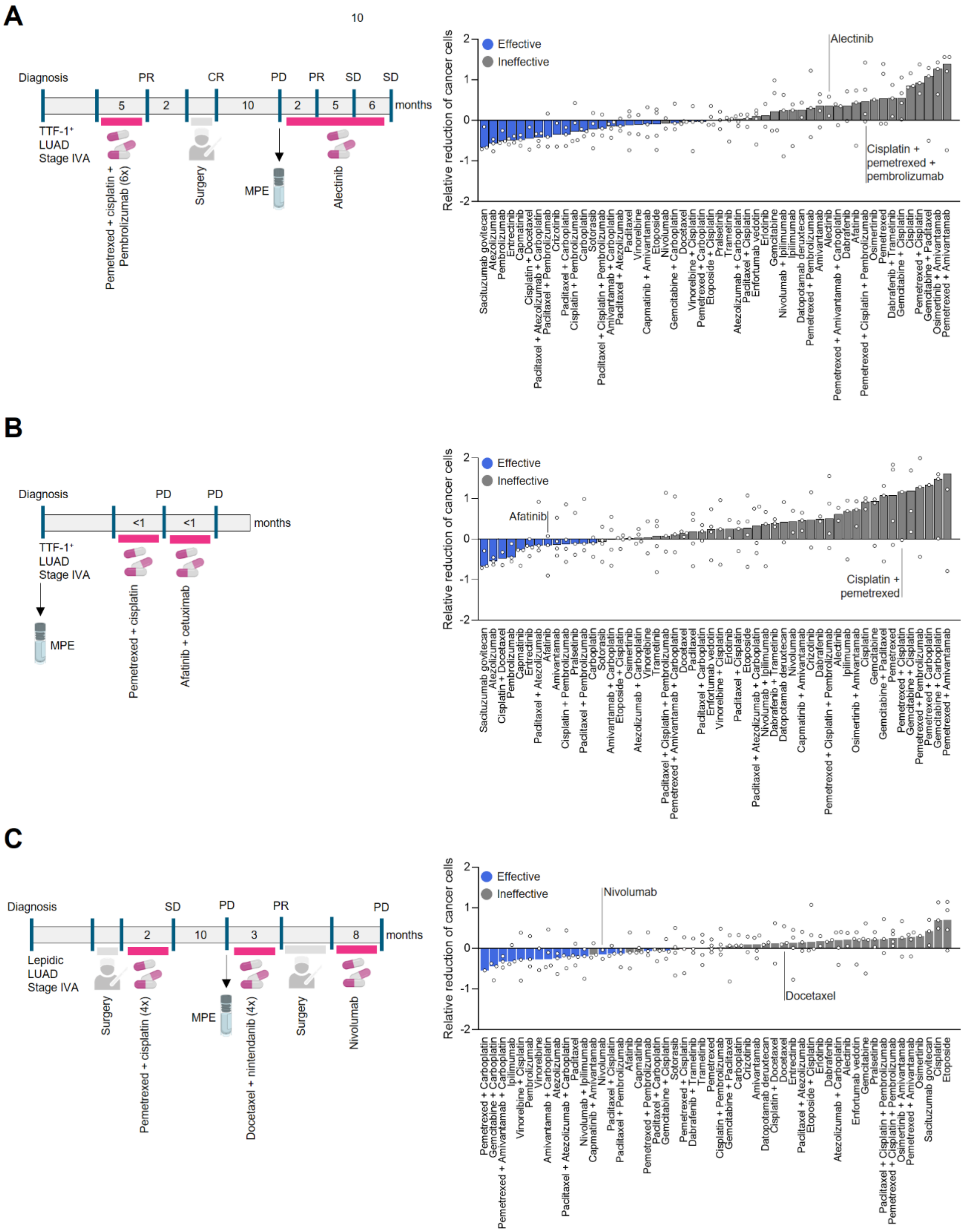


**Supplementary Fig. 3. Clinical courses and ex vivo drug response profiles of other patients.** (**A**) Clinical courses (left) and ranking of n = 52 guideline drugs based on their ability to reduce cancer cell counts (right) for patient P179. Bars represent median. Each dot represents one technical replicate well. Blue bars indicate effective drugs; treatments clinically administered to the patient are indicated in the plot. (**B**) As for (A), but for P206. (**C**) As for (A), but for P357.

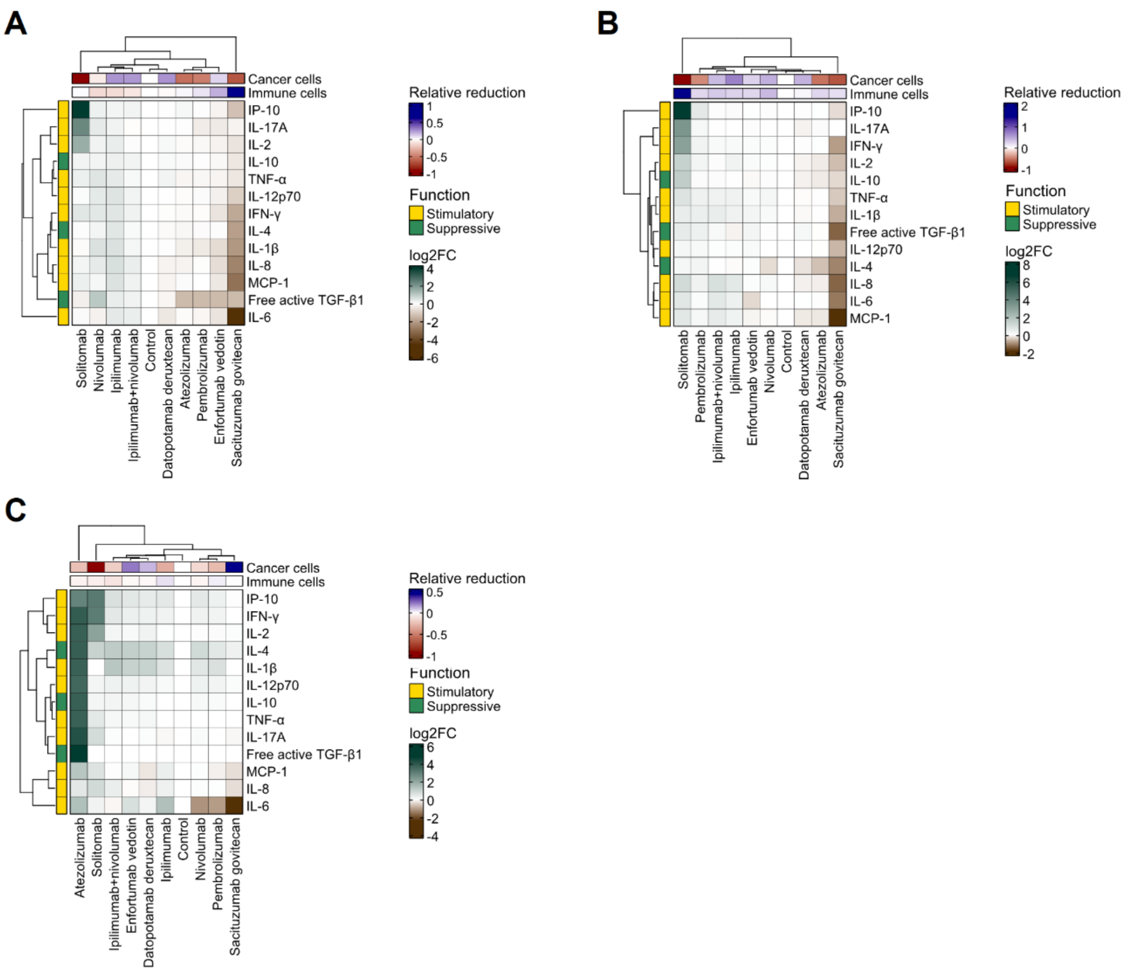


**Supplementary Fig. 4. Immunomodulatory profiles of additional MPE samples.** (**A**) Cytokine secretion matrix of patient P179 for n = 13 cytokines (rows) across immunomodulatory drugs (n = 9; columns), measured by multiplex cytokine assay. Scores (color scale) indicate Log2-fold changes of concentrations relative to isotype control. (**B**) As for (A), but for P206. (**C**) As for (A), but for P357.

### SUPPLEMENTARY TABLES

#### Supplementary Table 1. Baseline cytokine production of MPEs.

| Conc. (pg/mL) | P357 | P369 | P206 | P179 | P358 |
|---|---|---|---|---|---|
| IL-4 | 0 | 6.79 | 7.06 | 6.32 | 1.04 |
| IL-2 | 0.05 | 1.49 | 0.98 | 0.67 | 0.1 |
| IP-10 | 2.45 | 21.73 | 20.6 | 15.66 | 5.46 |
| IL-1β | 0 | 7.02 | 4.78 | 3.55 | 0 |
| TNF-α | 0 | 1.352 | 0.875 | 0.419 | 0 |
| MCP-1 | 10.751 | 2786.056 | 1357.911 | 642.723 | 324.054 |
| IL-17A | 0.37 | 0.7 | 0.52 | 0.73 | 0.19 |
| IL-6 | 13.025 | 184.756 | 502.07 | 1941.546 | 59.049 |
| IL-10 | 0 | 0.93 | 0.94 | 0.61 | 0 |
| IFN-γ | 0 | 8.296 | 6.683 | 4.352 | 0.368 |
| IL-12p70 | 0.16 | 3.21 | 2.75 | 1.86 | 0.36 |
| IL-8 | 28.05 | 4002.5 | 1872.01 | 777.89 | 153.28 |
| Free active TGF-β1 | 0 | 5.09 | 1.57 | 1.72 | 0 |

#### Supplementary Table 2. Drug panel reflecting treatment guideline of NSCLC.

| Drug | ID | Class | Status (NSCLC) | Concentration |
|---|---|---|---|---|
| Afatinib | CHEMBL1173655 | TKIs | Approved | 0,052 µM |
| Alectinib | CHEMBL1738797 | TKIs | Approved | 1,38 µM |
| Amivantamab | CHEMBL4297774 | Bispecific antibodies | Approved | 10 µg/mL |
| Atezolizumab | CHEMBL3707227 | ICIs | Approved | 10 µg/mL |
| Capmatinib | CHEMBL3188267 | TKIs | Approved | 3.67 µg/mL |
| Capmatinib + Amivantamab | N/A | Combination therapies | Approved | 3.67 µg/mL + 10 µg/mL |
| Carboplatin | CHEMBL1351 | Chemotherapies | Approved | 135 µM |
| Carboplatin + Amivantamab | N/A | Combination therapies | Approved | 135 µM + 10 µg/mL |
| Carboplatin + Atezolizumab | N/A | Combination therapies | Approved | 135 µM + 10 µg/mL |
| Cisplatin | CHEMBL11359 | Chemotherapies | Approved | 14,4 µM |
| Cisplatin + Pembrolizumab | N/A | Combination therapies | Approved | 14,4 µM + 10 µg/mL |
| Crizotinib | CHEMBL601719 | TKIs | Approved | 0,913 µM |
| Dabrafenib | CHEMBL2028663 | TKIs | Approved | 4,86 µM |
| Dabrafenib + Trametinib | N/A | Combination therapies | Approved | 4,86 µM + 0,021 µM |
| Datopotamab deruxtecan | N/A | ADCs | Approved | 10 µg/mL |
| Docetaxel | CHEMBL3545252 | Chemotherapies | Approved | 5,47 µM |
| Docetaxel + Cisplatin | N/A | Combination therapies | Approved | 5,47 µM + 14,4 µM |
| Enfortumab vedotin | CHEMBL3301589 | ADCs | Approved | 10 µg/mL |
| Entrectinib | CHEMBL1983268 | TKIs | Approved | 2,38 µM |
| Erlotinib | CHEMBL553 | TKIs | Approved | 3,15 µM |
| Etoposide | CHEMBL44657 | Chemotherapies | Approved | 33,4 µM |
| Etoposide + Cisplatin | N/A | Combination therapies | Approved | 33,4 µM + 14,4 µM |
| Gemcitabine | CHEMBL888 | Chemotherapies | Approved | 89,3 µM |
| Gemcitabine + Carboplatin | N/A | Combination therapies | Approved | 89,3 µM + 135 µM |
| Gemcitabine + Cisplatin | N/A | Combination therapies | Approved | 89,3 µM + 14,4 µM |
| Gemcitabine + Paclitaxel | N/A | Combination therapies | Approved | 89,3 µM + 4,27 µM |
| Ipilimumab | CHEMBL1789844 | ICIs | Approved | 10 µg/mL |
| Nivolumab | CHEMBL2108738 | ICIs | Approved | 10 µg/mL |
| Nivolumab + Ipilimumab | N/A | Combination therapies | Approved | 10 µg/mL + 10 µg/mL |
| Osimertinib | CHEMBL3353410 | TKIs | Approved | 0,126 µM |
| Osimertinib + Amivantamab | N/A | Combination therapies | Approved | 0,126 µM + 10 µg/mL |
| Paclitaxel | CHEMBL428647 | Chemotherapies | Approved | 4,27 µM |
| Paclitaxel + Atezolizumab | N/A | Combination therapies | Approved | 4,27 µM + 10 µg/mL |

| | | | | |
|---|---|---|---|---|
| Paclitaxel + Carboplatin | N/A | Combination therapies | Approved | 4,27 µM + 135 µM |
| Paclitaxel + Carboplatin + Atezolizumab | N/A | Combination therapies | Approved | 4,27 µM + 135 µM + 10 µg/mL |
| Paclitaxel + Cisplatin | N/A | Combination therapies | Approved | 4,27 µM + 14,4 µM |
| Paclitaxel + Cisplatin + Pembrolizumab | N/A | Combination therapies | Approved | 4,27 µM + 14,4 µM + 10 µg/mL |
| Paclitaxel + Pembrolizumab | N/A | Combination therapies | Approved | 4,27 µM + 10 µg/mL |
| Pembrolizumab | CHEMBL3137343 | ICIs | Approved | 10 µg/mL |
| Pemetrexed | CHEMBL225072 | Chemotherapies | Approved | 100 µM |
| Pemetrexed + Amivantamab | N/A | Combination therapies | Approved | 100 µM + 10 µg/mL |
| Pemetrexed + Carboplatin | N/A | Combination therapies | Approved | 100 µM + 135 µM |
| Pemetrexed + Carboplatin + Amivantamab | N/A | Combination therapies | Approved | 100 µM + 135 µM + 10 µg/mL |
| Pemetrexed + Cisplatin | N/A | Combination therapies | Approved | 100 µM + 14,4 µM |
| Pemetrexed + Cisplatin + Pembrolizumab | N/A | Combination therapies | Approved | 100 µM + 14,4 µM + 10 µg/mL |
| Pemetrexed + Pembrolizumab | N/A | Combination therapies | Approved | 100 µM + 10 µg/mL |
| Pralsetinib | CHEMBL4297597 | TKIs | Approved | 2,83 µg/mL |
| Sacituzumab govitecan | CHEMBL3545262 | ADCs | Investigational | 10 µg/mL |
| Solitomab | CHEMBL2109264 | Bispecific antibodies | Not approved | 10 µg/mL |
| Sotorasib | CHEMBL4535757 | TKIs | Approved | 7.50 µg/mL |
| Trametinib | CHEMBL2103875 | TKIs | Approved | 0,021 µM |
| Vinorelbine | CHEMBL553025 | Chemotherapies | Approved | 0,811 µM |
| Vinorelbine + Cisplatin | N/A | Combination therapies | Approved | 0,811 µM + 14,4 µM |

#### Supplementary Table 3. Drug panel for investigational ADCs.

| Drug | ID | Class | Status (NSCLC) | Concentration |
|---|---|---|---|---|
| Datopotamab | CHEMBL5314627 | Antibodies | Control | 1 µg/mL<br>5 µg/mL<br>50 µg/mL |
| Datopotamab deruxtecan | CHEMBL4297939 | ADCs | Approved (accelerated) | 1 µg/mL<br>5 µg/mL<br>50 µg/mL |
| DM4 | N/A | Payloads | Control | 1 µM<br>5 µM<br>50 µM |
| Dxd | N/A | Payloads | Control | 1 µM<br>5 µM<br>50 µM |
| Enfortumab | CHEMBL3301579 | Antibodies | Control | 1 µg/mL<br>5 µg/mL<br>50 µg/mL |
| Enfortumab vedotin | CHEMBL3301589 | ADCs | Investigational | 1 µg/mL<br>5 µg/mL<br>50 µg/mL |
| Ladiratuzumab | N/A | Antibodies | Control | 1 µg/mL<br>5 µg/mL<br>50 µg/mL |
| Ladiratuzumab vedotin | CHEMBL4298101 | ADCs | Investigational | 1 µg/mL<br>5 µg/mL<br>50 µg/mL |
| MMAE | N/A | Payloads | Control | 1 µM<br>5 µM<br>50 µM |
| Ozuriftamab | CHEMBL5095432 | Antibodies | Control | 1 µg/mL<br>5 µg/mL<br>50 µg/mL |

| | | | | |
|---|---|---|---|---|
| Ozuriftamab vedotin | CHEMBL5095285 | ADCs | Investigational | 1 µg/mL<br>5 µg/mL<br>50 µg/mL |
| Patritumab | CHEMBL2109406 | Antibodies | Control | 1 µg/mL<br>5 µg/mL<br>50 µg/mL |
| Patritumab deruxtecan | CHEMBL4594611 | ADCs | Not approved | 1 µg/mL<br>5 µg/mL<br>50 µg/mL |
| Telisotuzumab | CHEMBL3545419 | Antibodies | Control | 1 µg/mL<br>5 µg/mL<br>50 µg/mL |
| Telisotuzumab vedotin | CHEMBL3990032 | ADCs | Approved (accelerated) | 1 µg/mL<br>5 µg/mL<br>50 µg/mL |
| Trastuzumab | CHEMBL1201585 | Antibodies | Control | 1 µg/mL<br>5 µg/mL<br>50 µg/mL |
| Trastuzumab deruxtecan | CHEMBL4297844 | ADCs | Approved (accelerated) | 1 µg/mL<br>5 µg/mL<br>50 µg/mL |
| Tusamitamab | CHEMBL5095460 | Antibodies | Control | 1 µg/mL<br>5 µg/mL<br>50 µg/mL |
| Tusamitamab ravtansine | CHEMBL4298098 | ADCs | Not approved | 1 µg/mL<br>5 µg/mL<br>50 µg/mL |
